# Control of leaflet movement pattern by a novel PP2C phosphatase DLM1 in *Medicago truncatula*

**DOI:** 10.1101/2025.04.28.650909

**Authors:** Dian Zhou, Shiqi Guo, Wenjing Yang, Baolin Zhao, Weiyue Zhao, Shaoli Zhou, Yawen Mao, Hailong Zhang, Yuqi Fang, Liangliang He, Liling Yang, Changning Liu, Jianghua Chen, Quanzi Bai

## Abstract

Sessile plants exhibit diverse movement behaviors that have long intrigued the scientists. The legume plants display a rhythmic leaflet movement pattern characterized by horizontal opening during the day and vertical closure at night. However, the underlying mechanisms remain largely enigmatic. Here, we isolated a mutant designated as *dlm1* (*downward leaflet movement1*) from *Medicago truncatula* that displays leaflets downward opening during daytime while upward closure at night. Cellular analyses reveal that this aberrant phenotype correlates with abnormal volume changes of motor cells within the pulvinus. *DLM1* exhibits high expression level in the motor organ pulvinus and encodes an unreported nuclear-localized PP2C phosphatase. Notably, phylogenetic analysis demonstrates that species exhibiting rhythmic leaflet movements consistently retain DLM1 homologs, suggesting the possibility of its functionally conserved role in regulating leaflets movement pattern. Structural characterization reveals that DLM1 possesses both phosphatase and kinase domains. Functional complementation assays demonstrate that the phosphatase domain is necessary and sufficient for maintaining the leaflet movement pattern. Collectively, our work uncovers a novel PP2C protein that governs the leaflet movement, providing mechanistic insights into this intriguing plant behavior.

## Introduction

Scientists have long been interested in plant movement and the underlying molecular mechanisms (*Darwin, 1880*), such as the heliotropic movement of sunflowers (*Atamian et al., 2016*), the gravitropic response of plant roots to gravity (*Chen et al., 2023; Nishimura et al., 2023*) and the seismonastic movements of carnivorous plants capturing insects (*Procko et al., 2022; Scherzer et al., 2019*). The legume species exhibit a circadian leaf movement pattern with leaflets open horizontally during the day and close vertically at night. This movement behavior initiates upon day-night transition and is driven by the motor organ pulvinus, which is located at the base of leaflets. To date, key factors responsible for pulvinus development have been identified, including the transcription factor ELP1 (ELOGATED PETIOLULE1) (*Chen et al., 2012; Hagihara et al., 2022; Zhou et al., 2012*), the cyclin protein GmILPA1 (Increased Leaf Petiole Angle1), the F-box proteins MIO1 (MINI ORGAN1)/SLB1 (SMALL LEAF AND BUSHY1) (*Yin et al., 2020; Zhou et al., 2021*), as well as the plant hormones BR and auxin, all of which play a role in pulvinus-driven leaf movement process (*Bai et al., 2022; Kong et al., 2021; Zhao et al., 2020; Zhao et al., 2021*).

Additionally, it has been proposed that signal transduction influences the ion flow through pulvinus cells, resulting in the asymmetrical and reversible swelling/shrinking of motor cells (*Cote, 1995; Kim et al., 1993; Kong et al., 2020; Moran, 2007; Oikawa et al., 2018; Satter, 1975; Satter and Galston, 1971*), thereby triggering the leaflet moves between horizontal and vertical positions. When the adaxial extensor cells release K^+^, C1^-^ and water to shrink, meanwhile abaxial flexor cells take up K^+^, C1^-^ and water to swell, leaflets close. They open when the reverse ion movements and turgor changes occur (*Cote, 1995; Kim et al., 1993; Kong et al., 2020; Moran, 2007; Moshelion et al., 2002; Oikawa et al., 2018; Satter, 1975; Satter and Galston, 1971*). The leaflets maintain a horizontal position after the open process while a vertical position after the close process. However, how the legume plants maintain this special movement pattern, in which leaflets horizontally open during the day and close vertically at night, remains poorly understood.

Protein phosphorylation and de-phosphorylation are reversible reactions (*Bhaskara et al., 2019*). PP2C family proteins are the most abundant phosphatases in plants and involved in various de-phosphorylation-related biological processes (*Kerk et al., 2008*). The most typical functions include members of subfamily A, which participate in developmental processes and stress responses in response to the plant hormone ABA (*Komatsu et al., 2013; Miao et al., 2020*). Additionally, members in clade D are involved in the leaves and flowers development (*Song and Clark, 2005; Yu et al., 2003*), while members in clade B and C have been found to participate in the protein kinase signaling pathway (*Schweighofer et al., 2007; Smékalováet al., 2014*) and deactivate H^+^-ATPase (*Spartz et al., 2014*), respectively. Furthermore, PP2C family members in clade E-G have been demonstrated to participate in stress response (*Bhaskara et al., 2017; Ding et al., 2018; Guo et al., 2022; Liu et al., 2012*). Despite the extensive study of PP2C proteins belonging to several subfamilies, the functions of most other PP2C subfamily members are still unknown, especially in legume plants.

Here, we characterized the *DLM1* (*Downward Leaflet Movement1*) gene encoding an unreported PP2C subfamily L member PP2C75 to orchestrate the leaflet movement pattern in *M. truncatula*. In the daytime, the *dlm1* mutant pulvinus cells were sharply shrunk on the abaxial side while swelled on the adaxial side, which is associated with the downward leaf movement phenotype. We identified that DLM1 contains both phosphatase and kinase domains and are evolutionarily conserved across leaflet movement species. Additionally, the phosphatase domain of the DLM1 protein is essential for maintaining the leaflet movement pattern. Our study provides a novel insight into understanding the underlying mechanism of leaflet movement pattern maintenance as well as the function of PP2C proteins in legume plants.

## Results

### Isolation and characterization of the *dlm1* mutant in *M. truncatula*

To gain a better understanding of the molecular mechanisms underlying leaflet movement, we screened the *Tnt1* insertion population of *M. truncatula* to identify mutants with abnormal leaflet movement pattern through forward genetic approaches (*Tadege et al., 2008*). Under normal light-dark cycle conditions, the wild-type (R108) leaflets expand to maintain a horizontal position during the day and close upwards at night (Figure 1A and 1B). However, we identified a mutant with an abnormal leaf movement pattern where the leaflets cannot keep a horizontal position during the day and exhibit downward movement phenotype (Figure 1A). Therefore, we named *dlm1* (*downward leaflet movement 1*). Further phenotypic analysis found that the *dlm1* mutant leaflets close upwards at night, displaying a behavior similar to the wild-type (Figure 1B). Statistical analysis of lateral leaflets angles further confirmed that we have identified a *dlm1* mutant that exhibits an abnormal leaflet movement pattern, with leaflets opening downward during the day and closing upward at night (Figure 1C, Figure S1).

**Figure. 1.**
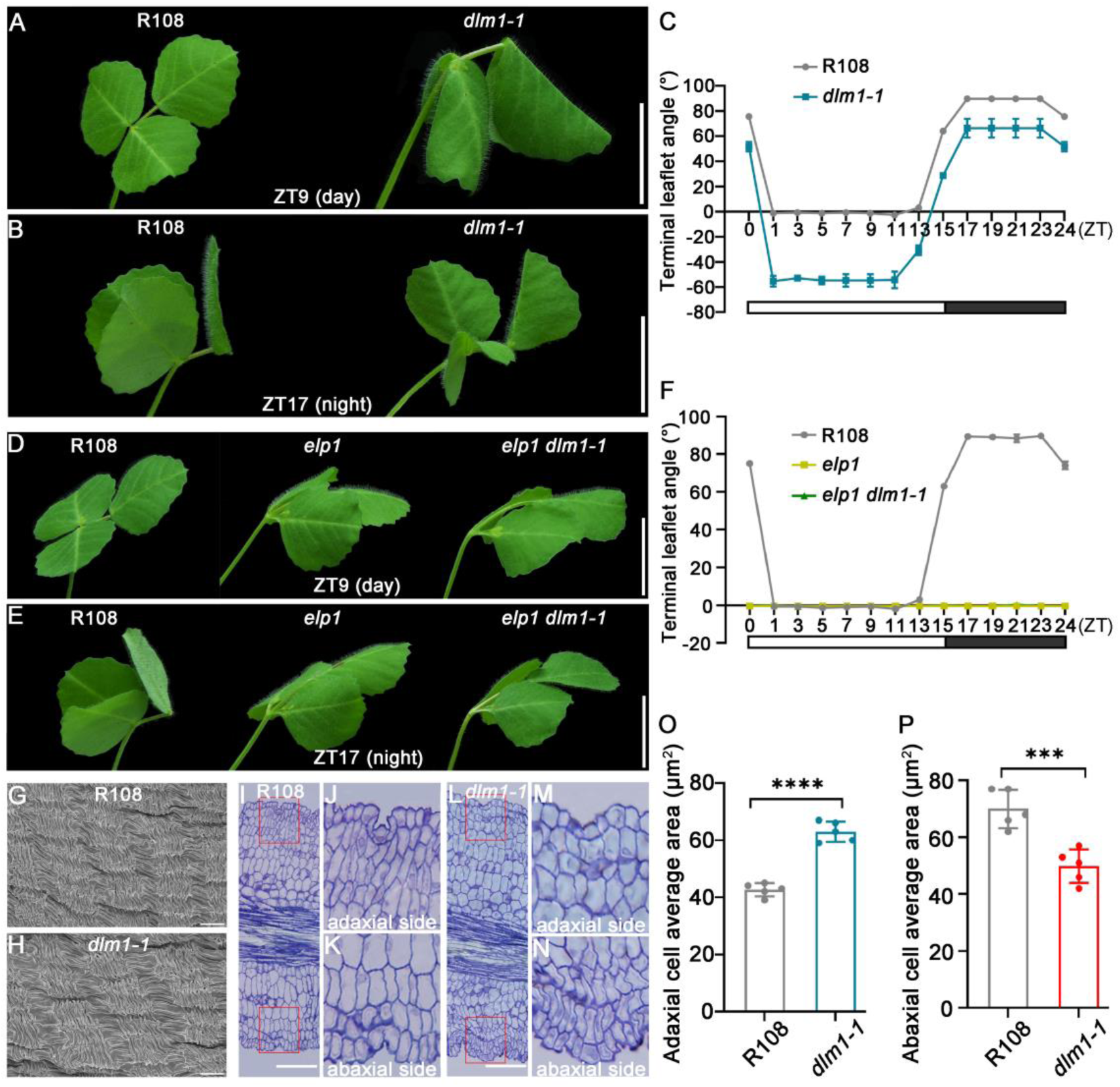
**The leaf movement phenotype of *dlm1* mutant in *M. truncatula*** (A, B) The leaflet movement phenotype of 6-week-old R108 (wild-type) (left) and *dlm1-1* mutant (right) at ZT9 (A) and ZT17 (B). Scale bars: 1 cm. (C) Statistics of terminal leaflet angles of WT and *dlm1-1* from ZT0 to ZT24 (ZT: zeitgeber). Data shown mean ± SD (n =3). (D, E) Genetic interactions between *elp1* and *dlm1-1* mutants. The leaflet movement phenotype of R108 (left), *elp1* (middle), and *elp1 dlm1-1* double mutant (right) at ZT9 (D) and ZT17 (E). Scale bars: 1 cm. (F) Statistics of terminal leaflet angles of WT, *elp1* and *elp1 dlm1-1* from ZT0 to ZT24 (ZT: zeitgeber). (G, H) SEM images of the epidermis of pulvini of 6-week-old R108 and *dlm1-1* plants. Scale bars: 10 µm. (I-N) Semi-thin sections of the mature pulvini structure of R108 and *dlm1-1* at ZT9. Scale bars: 50 µm. (O-P) Statistical analysis of data illustrates the average area of about four layers of adaxial and abaxial cells in the red box with five biological replicates, respectively at ZT9. (***, *P* < 0.001; ****, *P* < 0.0001; unpaired *t*-test).

Our previous studies have shown that *ELP1* determines the pulvinus organ identity in *M. truncatula*, loss of function of *ELP1* results in failing to fold leaflets due to loss of motor organs (*Chen et al., 2012*). To investigate whether the downward leaflet movement of *dlm1* is driven by the *ELP1*-dependent motor organ pulvinus, we generated *elp1 dlm1-1* double mutants. Phenotype analysis revealed that, similar to the *elp1* single mutant, the double mutant loses the pulvinus identity and leaflet movement behavior (Figures 1D-1F), indicating that the leaflet movement pattern of *dlm1* mutant is driven by the motor organ pulvinus. We then performed scanning electron microscopy (SEM) to analyze the epidermal structure of pulvini. The results showed that the surfaces of pulvini epidermal cells of *dlm1-1* were highly convoluted, giving the appearance of knitted wool, which is highly similar to the WT (Figures 1G and 1H).

Previous studies indicated that the reversible cell turgor-induced volume changes on the opposite side of pulvinus could drive leaf movement behavior. When the adaxial extensor cells release K^+^ and C1^-^ and shrink, meanwhile abaxial flexor cells take up K^+^ and C1^-^ and swell, leaflets close. They open when the reverse ion movements and turgor changes occur (*Cote, 1995; Kim et al., 1993; Oikawa et al., 2018*). To make clear the dynamic volume change of motor cells in *dlm1* mutant pulvinus, we used semi-thin sections analysis of pulvinus at ZT9 (zeitgeber time, 9 h after lights on) and ZT17, corresponding to the downward and upward leaflet movement state during the day and night, respectively. Results showed that both the WT and *dlm1-1* mutant displayed the normal pulvinus structure with layers of motor cells surrounding the central vascular bundle (Figures 1I and 1L). Further comparison analysis with the WT found that, at ZT9 the *dlm1-1* mutant motor cells of pulvini were sharply shrunk on the abaxial side while swelled on the adaxial side (Figures 1J-1K, 1M and 1N, 1O and 1P), which is consistent with the downward leaf movement phenotype. Reasonably, when compared at ZT17, both the motor cells of pulvini of WT and *dlm1* mutant were significantly shrunk on the adaxial side while swelled on the abaxial side, which is consistent with the upward leaf movement phenotype (Figure S2). Taken together, our results indicate that the dynamic volume change of motor cells mutant is associated with the leaflet movement pattern of *dlm1* mutant.

#### *DLM1* encodes a PP2C phosphatase to control leaflet movement

To identify the gene responsible for the *dlm1-1* mutant phenotype, we first backcrossed to the wild-type (WT, R108 ecotype) to purify the *Tnt1* insertion background. The F2 offspring of self-pollinated F1 plants showed a segregation ratio of 3:1, indicating that *dlm1-1* is caused by a single recessive nuclear mutation. Subsequently, we collected an equal amount of leaflet tissue from eighteen *dlm1-1* mutants from the BC1F2 population for whole genome resequencing at 20×coverage. This analysis identified a total of eighteen homozygous *Tnt1* insertions in the *dlm1-1* genome (Table S1). Genetic linkage analysis revealed that the mutation may be caused by the reverse homozygous insertion of *Tnt1* into the first exon of the *DLM1* (*Medtr6g087000/MtrunA17_Chr6g0486661*) gene (Figure 2A, Figure S3). The amino acid sequence alignment results indicated that *DLM1* encodes a protein phosphatase homologous to the *Arabidopsis* PP2C75 (AT2G20050), which contains an N-terminal phosphatase domain and C-terminal kinase domain (Figures 2B, Figure S4).

**Figure. 2.**
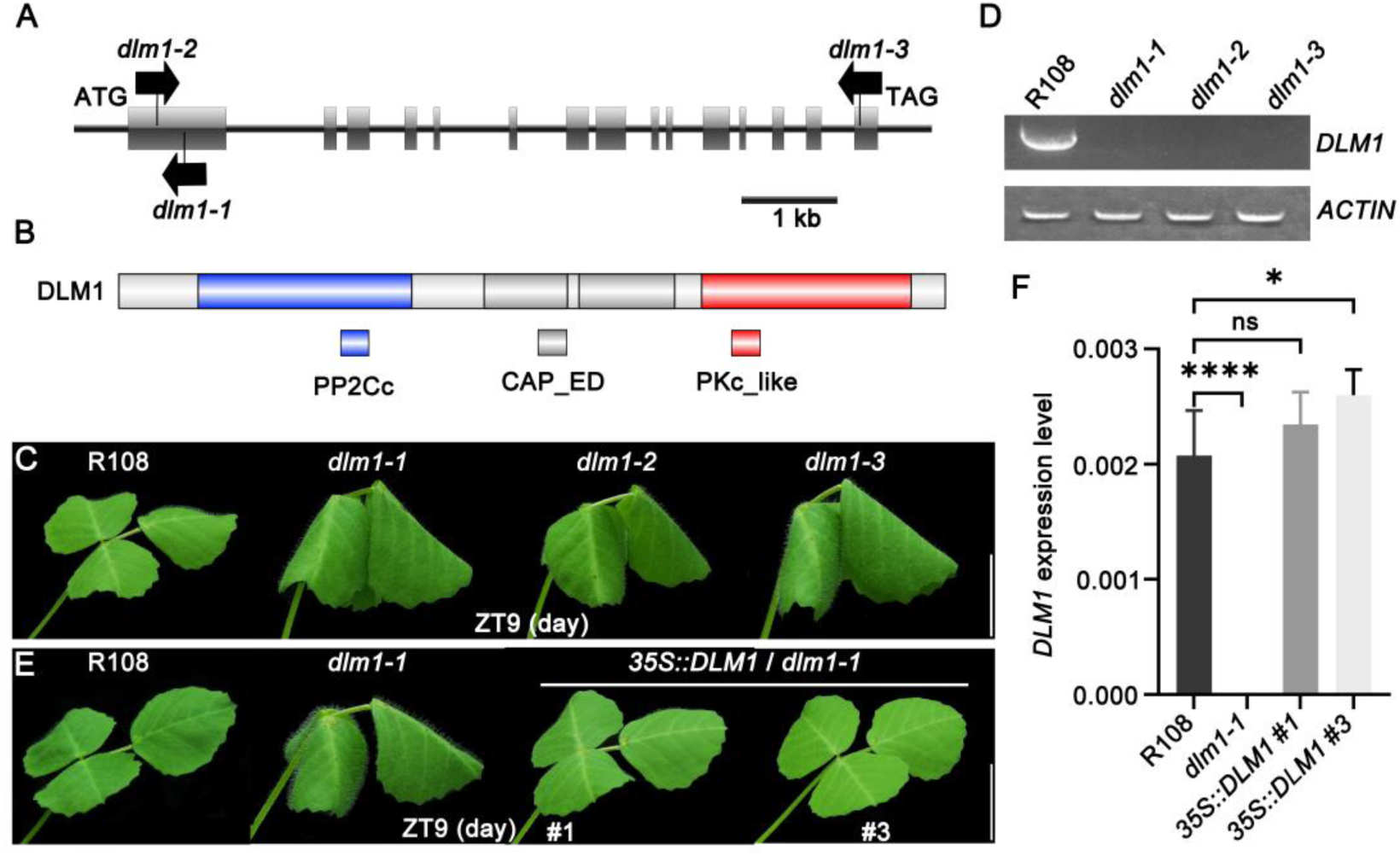
**Molecular cloning of the *DLM1*** (A) The gene structure of *DLM1* and *Tnt1* insertion sites of three different alleles. The start codon (ATG) and stop codon (TAG) are shown, black lines represent introns and the grey boxes represent exons among them. The arrows indicate the orientation of *Tnt1* insertions. (B) Analysis of DLM1 protein structural domain. The blue box represent PP2C phosphatase domain, The red box mean PKc_like kinase domain, and the grey box represent CAP_ED domain, which is the effector domain of the CAP family of transcription factors. (C) The phenotype of *dlm1* alleles. The leaflet movement phenotype of 6-week-old R108 (left) and *dlm1-1*, *dlm1-2,* and *dlm1-3* mutants (right) at ZT9. Scale bars: 1 cm. (D) RT-PCR analysis of *DLM1* expression in pulvini of R108 and three *dlm1* mutant alleles. (E) Genetic complementation of *dlm1*. The representative *dlm1-1* mutant was transformed with *35S::DLM1*. The two rescued mutant plants (*35S::DLM1*#1 and *35S::DLM1*#3) are shown during the day. Scale bars: 1 cm. (F) The expression level of *DLM1* by RT-qPCR in pulvini of R108, *dlm1-1,* and two independent *35S::DLM1*/*dlm1-1* transgenic lines. Asterisks indicate differences between R108, *dlm1-1,* and two transgenic lines. (*, *P* < 0.05; ****, *P* < 0.0001; ns: no significant difference, one-way ANOVA.)

In order to confirm the candidate gene, we further screened the public mutation database and isolated two other *Tnt1* inserted alleles, *dlm1-2* and *dlm1-3*, both exhibited downward leaflet movement at ZT9 and upward leaflet movement at ZT17, displaying movement pattern similar to *dlm1-1* (Figures 2A and 2C, Figure S5A). RT-PCR showed undetectable transcript levels of *Medtr6g087000* in the three *dlm1* mutant alleles, indicating successful knockout of candidate gene in the *dlm1* alleles (Figure 2D). Then, we introduced *35S::DLM1* into the *dlm1-1* mutants and generated two independent transgenic lines. Phenotype and RT-qPCR analyses results indicated that overexpression of *DLM1* completely restored the abnormal leaf movement phenotype of *dlm1-1* mutant (Figures 2E and 2F, Figure S5B). Collectively, these results suggest that the gene *Medtr6g087000/MtrunA17_Chr6g0486661* is responsible for the *dlm1* downward leaf movement phenotype.

#### Expression pattern of *DLM1* and subcellular localization of its encoding protein

To better investigate the role of *DLM1* in regulating leaf movement in *M. truncatula*, we characterized its expression pattern. RT-qPCR results demonstrated that *DLM1* is highly expressed in the pulvinus organ, with the highest levels found in pulvinus (Figure 3A). To further examine the temporal and spatial expression patterns of *DLM1* in pulvinus tissue, we then generated *pDLM1::GUS* transgenic plants in the R108 ecotype. GUS staining assays indicated that the signal was predominantly enriched in the pulvinus and leaves (Figures 3B and 3C). Both of the RT-qPCR and GUS staining results suggest that *DLM1* is highly expressed in pulvinus, which is consistent with its function in leaf movement behavior.

**Figure. 3.**
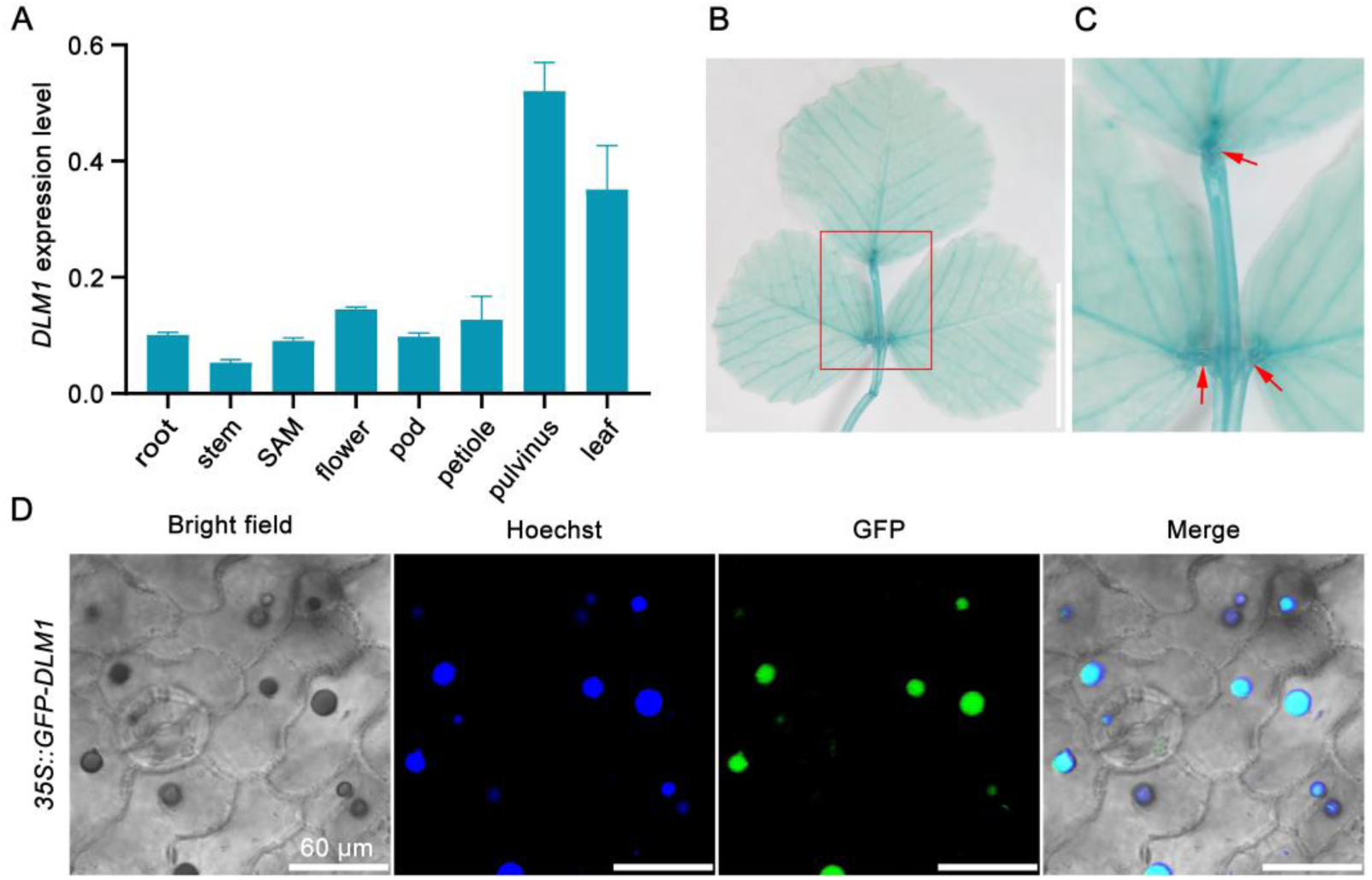
**Expression pattern of *DLM1* and subcellular localization of its encoding protein** (A) The expression level of *DLM1* in different tissues in *M. truncatula*. (B, C) GUS staining of compound leaves of 6-week-old *pDLM1::GUS* transgenic plants. The red box indicates the enlarged area, and the red arrows indicate the pulvinus area. Scale bar: 1 cm. (D) Subcellular localization analysis of DLM1 with *35S::GFP-DLM1* vector in *Arabidopsis thaliana* (Col-0). Scale bars: 60 µm.

To explore the role of DLM1 at the cellular level, we performed subcellular localization analysis by using *35S::GFP-DLM1* transformed *Arabidopsis thaliana*. The results showed that the GFP signal was specifically detected in the nucleus, indicating that the DLM1 protein is localized to the nucleus. This finding further suggests that DLM1 functions within the nucleus (Figure 3D).

#### Phylogenetic analysis of DLM1 and its homologs

Traditional studies identified that PP2C family proteins function as a major phosphatase group and participate in many key biological processes in plants (*Kerk et al., 2008*). Phylogenetic analysis of DLM1 in the representative species including spore plants (*Chlamydomonas reinhardtii*, *Micromonas pusilla*, *Marchantia polymorpha*, *Physcomitrium patens*, *Diphasiastrum complanatum*, *Ceratopteris richardii*, *Marsilea vestita*), gymnosperm (*Pinus taeda*, *Ginkgo biloba*) and angiosperm (*Amborella trichopoda*, *Oryza sativa*, *Arabidopsis thaliana*, *Solanum lycopersicum*, *Averrhoa carambola*, *Euphorbia peplus*, *Sauropus androgynus*, *Cucumis sativus*, *Pisum sativum*, *Medicago truncatula*, *Lotus japonicas*) indicated that *DLM1* encodes a protein phosphatase belonging to the L subfamily of the PP2C protein phosphatase family (Figures 4A and B). To our knowledge, DLM1 is the first member of this subfamily which has been functionally characterized in plants to date. Additionally, phylogenetic analysis revealed that DLM1 homologs are widely conserved across species, all containing both phosphatase and kinase domains (Figure 4C). Notably, species displaying pulvinus-driven leaflets movement behavior including *Medicago truncatula*, *Pisum sativum*, *Lotus japonicas, Sauropus androgynus*, *Averrhoa carambola* and *Marsilea vestita* consistently retain DLM1 homologs (Figure 4C), indicating that DLM1 may have a functionally conserved role in regulating leaflets movement patterns. Together, these results suggest that DLM1 represents a functionally uncharacterized PP2C family member that may be evolutionarily conserved to regulate leaflet movement patterns.

**Figure. 4.**
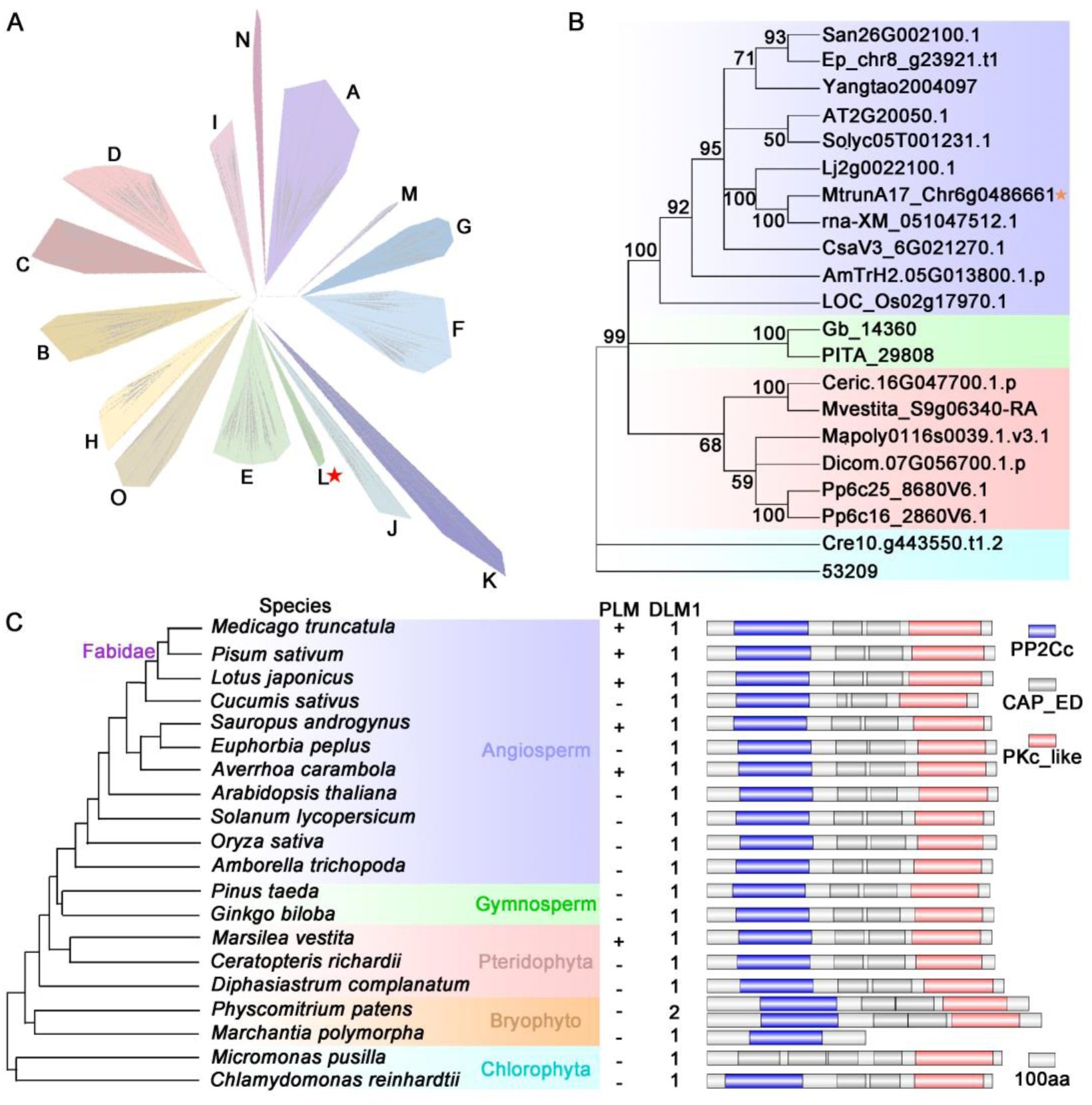
**Phylogenetic analysis of DLM1 and its homologs** (A) Neighbor-joining tree of PP2C family proteins from 20 species. Each colored block represents a subfamily of PP2C proteins from multiple species. The red pentagram represents the PP2C L subfamily where DLM1 is located. (B) Magnified view of PP2C L subfamily from (A). DLM1 and its homologs present in species including *Sauropus androgynus* (San), *Euphorbia peplus* (Ep), *Averrhoa carambola* (Yangtao), *Arabidopsis thaliana* (AT), *Solanum lycopersicum* (Solyc), *Lotus japonicus* (Lj), *Medicago truncatula* (Mt), *Pisum sativum*, *Cucumis sativus* (Csa), *Amborella trichopoda* (AmTr), *Oryza sativa* (Os), *Ginkgo biloba* (Gb), *Pinus taeda* (PITA), *Ceratopteris richardii* (Ceric), *Marsilea vestita* (Mvestita), *Marchantia polymorpha* (Mapoly), *Diphasiastrum complanatum* (Dicom), *Physcomitrium patens* (Pp), *Chlamydomonas reinhardtii* (Cre), *Micromonas pusilla*. The orange pentagram indicates the DLM1 position. The purple, green, pink and blue boxes represent the classification of angiosperms, gymnosperms and spore plants, respectively. (C) The presence of leaf movement species, DLM1 homologs and their domain architecture.

#### DLM1 employs its N-terminal phosphorylase domain to regulate leaflet movement pattern

Protein phosphorylation and de-phosphorylation are reversible reactions, which have been found to be involved in numerous biological processes in eukaryotes. Protein kinases (PKs) and phosphatases (PPs) function as the “writer” and “eraser” to mediate these reversible events (*Bhaskara et al., 2019*), respectively. Intriguingly, here we identified DLM1 protein harbors both protein phosphatase and protein kinase domain, a structure that hasn’t been reported before in plants. Based on this finding, we supposed that both the N-terminal phosphorylase domain and the C-terminal kinase domain were essential to regulate the leaflet movement pattern. To verify our hypothesis, we constructed several truncated *DLM1* sequences driven by 35S promoter (Figure 5A), respectively, and then introduced them into the *dlm1-1* mutant plants to generate different transgenic lines to examine if they could rescue the mutant downward leaflet movement phenotype. However, the results are much different from our initial expectations. Results showed that both the *35S::DLM1* (*DLM1* whole coding sequence) and *35S::N-DLM1* (with deletion of C-terminal Ser/Thr kinase domain sequence) could rescue the downward leaflet movement, while *35S::C-DLM1* (with deletion of N-terminal PP2C protein phosphorylase domain sequence) could not (Figures 5B-5E). Taken together, our truncated complementation experiments revealed that the N-terminal phosphorylase domain is essential and sufficient to maintain the leaflet in a horizontally open state during the day, thus regulating the leaflet movement pattern in *M. truncatula*.

**Figure. 5.**
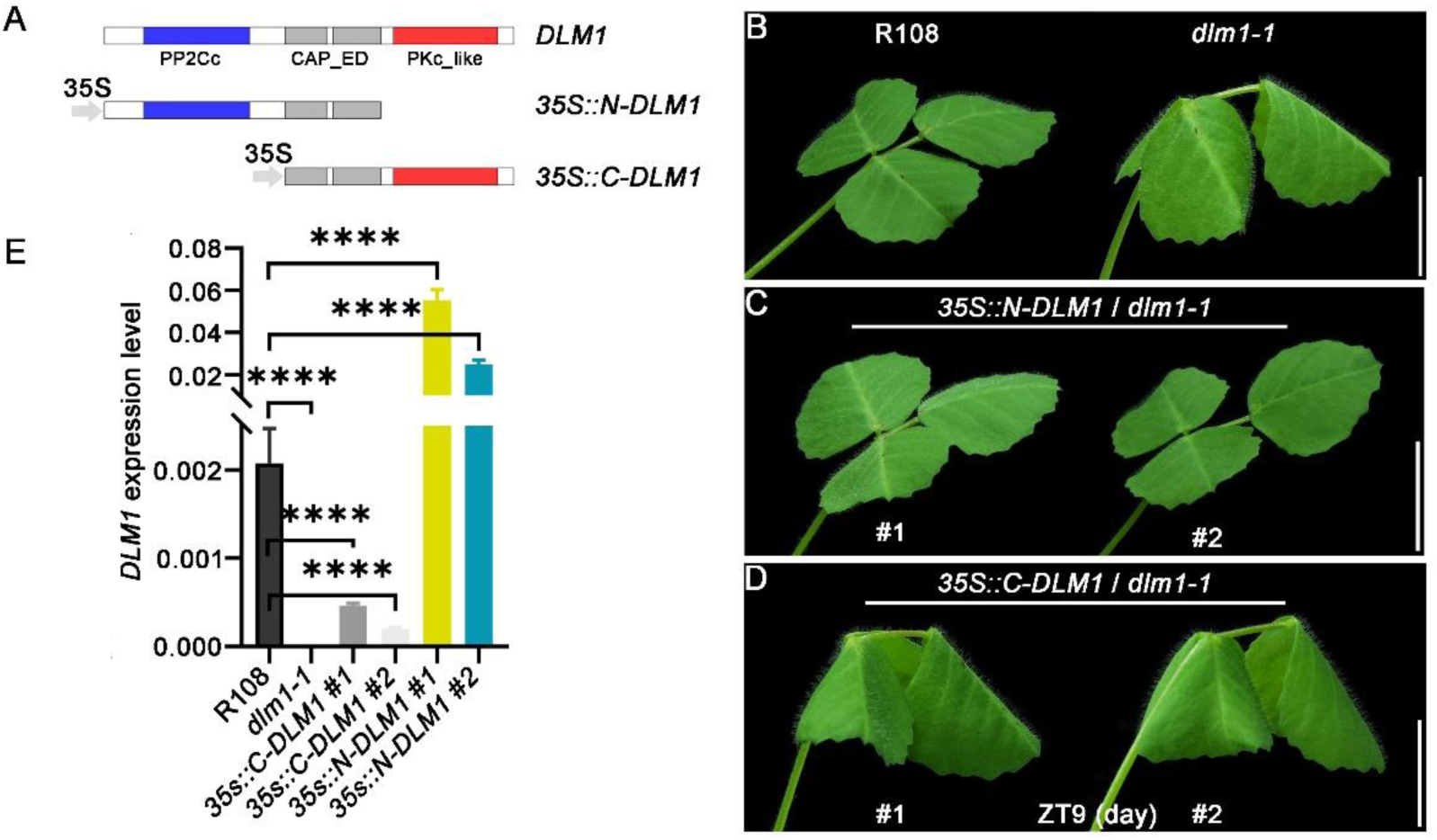
**Protein domain and functional analysis of DLM1** (A) The Schematic representation of constructs used for transformation. (B-D) Transgenic truncated complementation of *dlm1-1*. (B) The phenotype of R108 and *dlm1* plants during the day (top). (C) The two rescued mutant plants of *35S::N-DLM1*#1 and *35S::N-DLM1*#2 are shown at ZT9 (middle). (D) While the *35S::C-DLM1*#1 and *35S::C-DLM1*#2 two lines are also shown at ZT9 (below). Scale bars: 1 cm. (E) RT-qPCR analyses the expression of truncated transgenic plants. ****, *P* < 0.0001, one-way ANOVA.

## Discussion

The legume plants utilize the motor organ pulvinus to drive their leaflets open horizontally and close vertically upon day-night transition. Key factors including ELP1 (*Chen et al., 2012; Hagihara et al., 2022; Zhou et al., 2012*), MIO1/SLB1 (*Zhou et al., 2021*), ILPA1 (*Gao et al., 2017*), phytohormones (*Bai et al., 2022; Kong et al., 2021; Zhao et al., 2020; Zhao et al., 2021*), light (*Kim et al., 1993*), circadian clock (*Kim et al., 1993; Kong et al., 2020; Oikawa et al., 2018; Satter, 1975*), membrane ion/aquaporin transporters (*Kim et al., 1993; Moshelion et al., 2002; Oikawa et al., 2018*) have been identified to orchestrate the open or close process. However, the mechanisms by which the horizontal-vertical leaflet movement pattern is maintained remain poorly understood. Through forward genetics, this study identified the *dlm1* mutant exhibits a rhythmic downward-upward leaflet movement pattern. Additionally, when crossed with the pulvinus-deficient mutant *elp1* to generate *elp1 dlm1-1* double mutant, all downward and upward leaflet movement phenotypes were absent (Figures 1D-1F). Thus, we identified a key factor in maintaining the leaflet movement pattern through pulvinus.

During the leaflet movement process, the flow of K^+^ and C1^-^ through the membrane of pulvinus cells result in swell or shrink of the opposite cells, thereby driving the opening or closing process of the leaflet (*Kim et al., 1993; Oikawa et al., 2018*). This research found the mutation of DLM1 causes the shrinkage of the abaxial cells and subsequently drives the leaflet to move downward (Figures 1I-1N). To uncover the mechanism by which DLM1 regulates the volume changes of pulvinus abaxial cells, we analyzed the downstream biological processes affected by DLM1 at the transcriptional level. A total of 232 differentially expressed genes (DEGs), with 82 upregulated and 150 downregulated, were found in *dlm1-1* mutant pulvinus compared to wild-type R108 (Figure S6A). Gene Ontology (GO) enrichment analyses of these DEGs were significantly enriched in processes such as cell wall organization or biogenesis, and response to stimulus (Figure S6B). These data indicated the role of DLM1 in affecting the expression of many genes including cell wall-related genes. To further figure out the molecular mechanisms underlying leaflet movement, further studies are needed to investigate the relationship between DLM1 and the activity of ion channel proteins activity in pulvinus cells. Moreover, the abaxial cells in the *dlm1* mutant do not consistently maintain a contracted state and can also drive the leaflets to close upward through cell swelling (Figures 1A-1F), suggesting a complex regulatory mechanism underlying the dynamic volume changes of the pulvinus cells.

The phosphorylation and de-phosphorylation of biomolecules depend on protein phosphatases and kinases within living organisms (*Novak et al., 2010; Sojka et al., 2024*). As the largest family of protein phosphatases within plants, previous studies have reported that PP2C family proteins function by removing phosphate modifications from substrates through their phosphatase domains (*Ma et al., 2009b; Shi, 2009*). The most typical functions include members of subfamily A, which participate in developmental processes and stress responses in response to the plant hormone ABA (*Komatsu et al., 2013; Komatsu et al., 2020; Ma et al., 2009a*). This study identified a new phosphatase in *M. truncatula*, PP2C75, which belongs to the L subfamily of the PP2C protein phosphatase family and has not been functionally reported before. Although functionally uncharacterized previously, we observed the persistent presence of DLM1 in leaf-movement species spanning ferns to advanced angiosperms, implying possible functional conservation in leaflet movement control. Further investigations are needed to confirm this hypothesis.

Additionally, functional complementation assays demonstrate that the phosphatase domain is both necessary and sufficient for maintaining the leaflet movement pattern, further supporting the importance of phosphorylation signaling in regulating this process. These findings suggest that other phosphatases or kinases may also participate in this regulatory network, providing a direction for future research. Although the role of the phosphatase domain in *Medicago truncatula* leaflet movement has been established, the function of the kinase domain and the biological significance of homologous proteins in other species remain to be elucidated. Further investigation of these aspects will expand our understanding of PP2C functions and phosphorylation-dependent modifications in plant movement.

## Materials and Methods

### Plant materials and growth conditions

The *dlm1-1* mutants (NF7737) in *M. truncatula* were isolated and collected from the *Tnt1* retrotransposon tagged mutant collections by a forward-genetics approach as previously reported.(*Tadege et al., 2008*) As for the *dlm1-2* (NF1438) and *dlm1-3* (NF14116) plant materials were requested from the Noble Research Institute Mutant collections (https://medicago-mutant.dasnr.okstate.edu/mutant/index.php), and all plants used in these studies are all R108 ecotype. Plants grown in a greenhouse under such controlled conditions: 15 h/9 h day/night cycle, 70% humidity, 150 μmol/m2 /s light intensity, and 23 ℃/ 20 ℃ day/night temperature.

### Whole genome resequencing

The *dlm1-1* mutant population was backcrossed with the wild-type R108 ecotype and then bred for two generations, here 8-week-old *dlm1-1* and the wild-type-like heterozygous plants were sampled with three biological repetitions and the genomic DNA of was extracted by a Plant Genomic DNA Kit (Tiangen, Beijing, China) following the instructions. The DNA samples were used for genome resequencing at 20x coverage. The insertion sites of the *Tnt1* retrotransposon in the genome of *M. truncatul*a were analyzed in Table S1.

### Semi-thin sections

Pulvini from wild-type and mutant plants about eight weeks old were removed and fixed in a formalin-acetoalcohol (FAA) solution. Following ethanol gradient dehydration, semi-thin sections were cut on a Leica UC7 using ethylene glycol methacrylate (GMA) as the penetrant. They were stained with 0.05% toluidine blue for 10 to 20 s and visualized under an Olympus BX63 microscope to observe the histological characteristics of the pulvini.

### Quantification and statistical analysis

The semi-thin slice images were quantified using ImageJ. All statistical data were analyzed by GraphPad Prism version 8. The P-values were calculated by the ordinary ANOVA test one-way ANOVA, or using unpaired t test.

### Scanning electron microscopy (SEM)

Mature pulvini from eight-week-old dlm1-1 and R108 were collected and fixed in a 3% glutaraldehyde fixation solution (5% acetic acid, 5% formaldehyde, 45% ethanol). The tissues were washed, dehydrated, and dried at Kunming Botanical Institute. Then, they were coated with gold, and examined under a Sigma 300 microscope (Zeiss).

### Plasmid and transgenic plants construction

The coding sequence of *DLM1* was amplified by R108 pulvinus cDNA. The samples applied to DLM1, N-DLM1, and C-DLM1, were cloned into *E. coli* TOP10 strains and determined by Sanger sequencing, then recombined into vector *pCAMBIA3301-vocGFP-MP* to construct *35S::DLM1*, *35S::GFP-DLM1*, *35S::N-DLM1* and *35S::C-DLM1*, respectively. The constructs of *35S::DLM1*, *35S::N-DLM1,* and *35S::C-DLM1* were transformed into *dlm1-1* mutants via Agrobacterium-mediated transformation. For *DLM1* promoter activity analysis, the promoter sequence was fused to the *pCAMBIA3301* vector. The primer sequences are listed in Table S2.

### Protein sequence alignment and phylogenetic analysis

The amino acid sequences were sourced from Phytozome v13 (https://phytozome-next.jgi.doe.gov/). Multiple sequences were aligned using ClustalX software. Phylogenetic trees were constructed using the neighbor-joining method implemented in IQ-TREE2 (JTT+R9 substitution model, with 1000 SH-aLRT replicates and 1000 ultrafast bootstrap replicates). Amino acid domains were predicted using the National Center for Biotechnology Information (https://www.ncbi.nlm.nih.gov/Structure/cdd/wrpsb.cgi). The resulting tree will be edited and enhanced using iTOL (https://itol.embl.de/).

### Subcellular localization

The *35S::GFP-DLM1* was introduced into Agrobacterium (Agrobacterium tumefaciens) strain EHA105 and transformed into Arabidopsis (Arabidopsis thaliana, Col-0) using the floral dip method. The homozygous T3 seedlings were used for GFP fluorescence imaging by using laser confocal microscope.

### GUS Staining

Different maturity levels complete leaves of *pDLM1::GUS* line were taken and soaked in the GUS dye solution by using a GUS stain kit for 37℃ overnight. The samples were soaked and washed with 75% alcohol and observed under the stereoscopic microscope.

### Transcriptome analysis

Quality control of both raw RNA-seq and sRNA-seq reads was performed by using FastQC (v0.12.1). Subsequently, transcriptome analysis was carried out with the HISAT2 pipeline (*Pertea et al., 2016*). The *Medicago truncatula* genome version utilized was Mt4.0v1, obtained from Phytozome v13 (https://phytozome-next.jgi.doe.gov/). Differentially expressed genes were identified using GFOLD with default parameters (*Feng et al., 2012*).

### Quantitative real-time PCR (RT-qPCR)

RNA was extracted from plant tissues using TransZol (TransGen, Beijing, China), 2 µg RNA sample was transcribed into cDNA using the TransScript II One-Step gDNA Removal kit (Takara, Shiga, Japan). RT-qPCR was performed in three biological replicates using TransStart Tip Green qPCR SuperMix (TransGen) on a LightCylcer 480 device (Roche, Basel, Switzerland). Data are mean ± SD (n = 4) with three biological replicates, and normalized the reference gene *MtActin*. All the RT-qPCR products were examined by electrophoresis. The primer sequences are listed in Table S2.

## Acknowledgments

The authors would like to thank Zhijia Gu (KIB, CAS) for SEM technical support, we also thank the provision of *M. truncatula Tnt1* insertion lines by the Noble Research Institute. This work is supported by the National Natural Science Foundation of China grants (32170839, 32200290, 32170360, 32470334, U2102222 and 32070204), Youth Innovation Promotion Association CAS (2021395), the Yunnan Revitalization Talent Support Program (XDYC-QNRC-2022-0179) and the Natural Science Foundation of Yunnan Province of China (202401AT070258).

## Author Contributions

B.Q.Z. and C.J.H. designed research. Z.D., G.S.Q., Y.W.J. and B.Q.Z. performed experiments. All authors analyzed and/or interpreted the data. B.Q.Z., Z.D. and C.J.H. wrote the manuscript with help from co-authors.

## Supplementary data

**Figure. S1.**
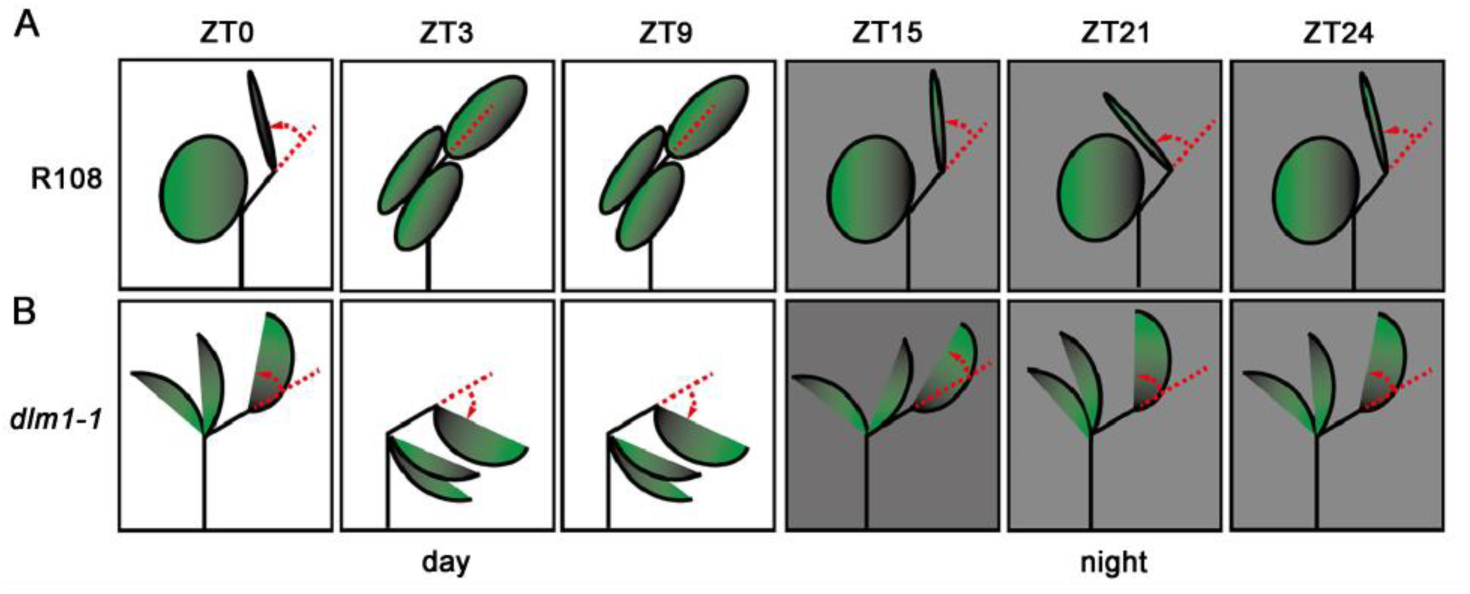
**Schematic diagram of terminal leaflet movement angle.** (A-B) Schematic diagram of measuring the angles of the terminal leaflets from daytime to nighttime for R108 (A) and *dlm1-1* (B). Using the leaf axis as a reference, the angle between the terminal leaflet and the leaf axis represents the leaf movement state for measurement.

**Figure. S2.**
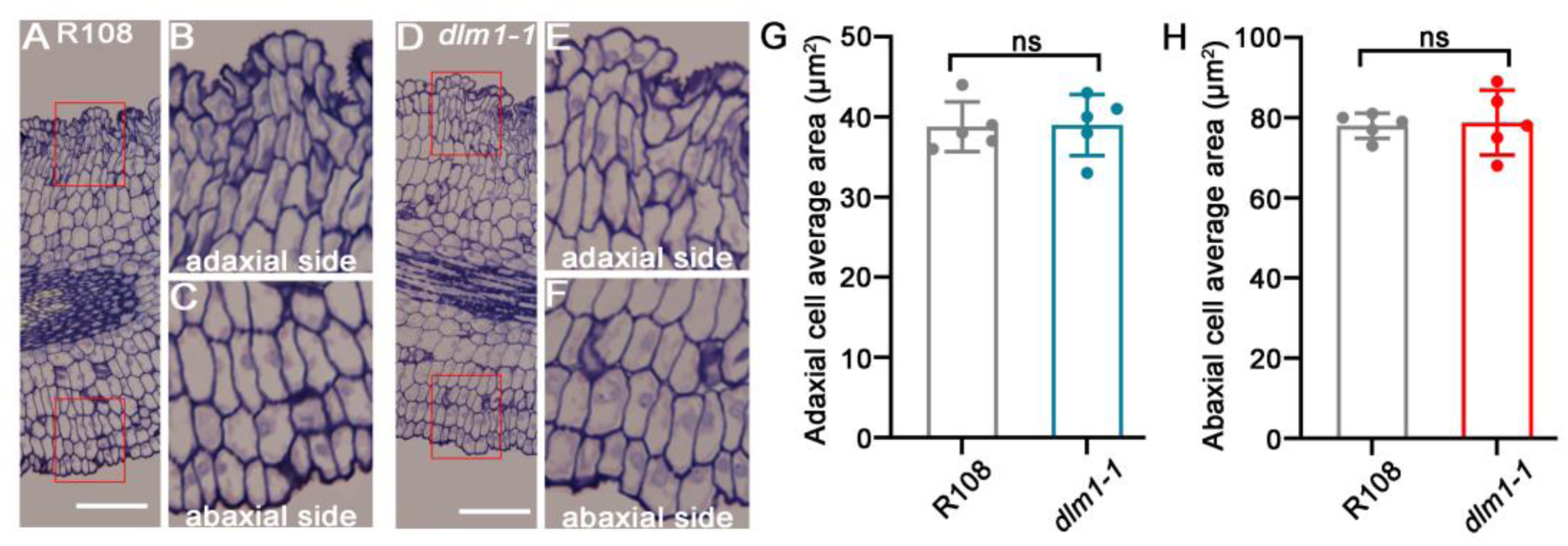
**Semi-thin sections of R108 and *dlm1-1* at ZT17.** (A-C) The Semi-thin sections of R108 at ZT17. (D-F) The Semi-thin sections of *dlm1-1* at ZT17 (ZT: zeitgeber). The red box shows the enlarged area. Scale bars:50 µm. (G-H) Statistical analysis of data shows the average area of approximately four layers of adaxial and abaxial cells in the red box with five biological replicates, respectively at ZT17. (ns: no significant difference; unpaired *t*-test.)

**Figure. S3.**
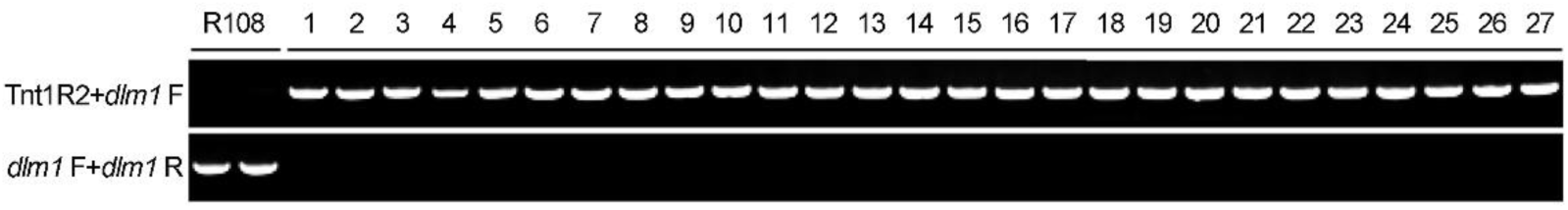
**Genetic linkage analysis of *dlm1*.** The twenty-seven BC1F2 population mutants of *dlm1* were used for genetic linkage analysis, and all are homozygous in *DLM1*, *dlm1*-F/R is a pair of specific primers for *DLM1* genomic sequence, *Tnt1* R2 is the retrotransponson *Tnt1* specific primer. The R108 is a control.

**Figure. S4.**
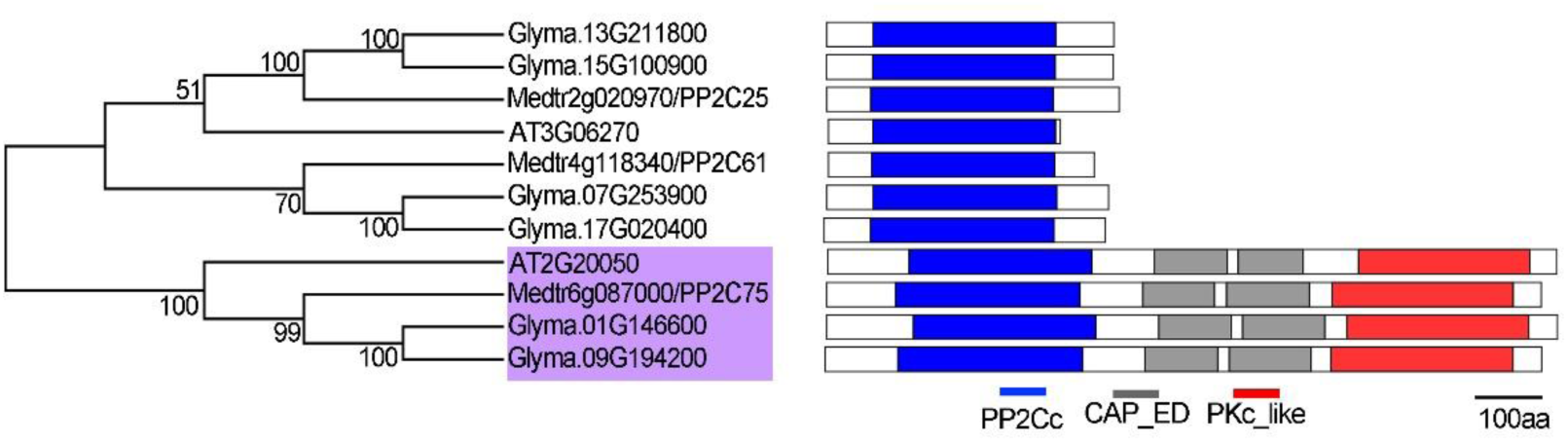
**Analysis of PP2C L subfamily conserved domains in *Medicago truncatula*, *Arabidopsis thaliana* and *Glycine max*.** The blue boxes represent PP2C phosphatase domain, grey boxes represent CAP_ED domain, and the red boxes represent PKc_like kinase domain.

**Figure. S5.**
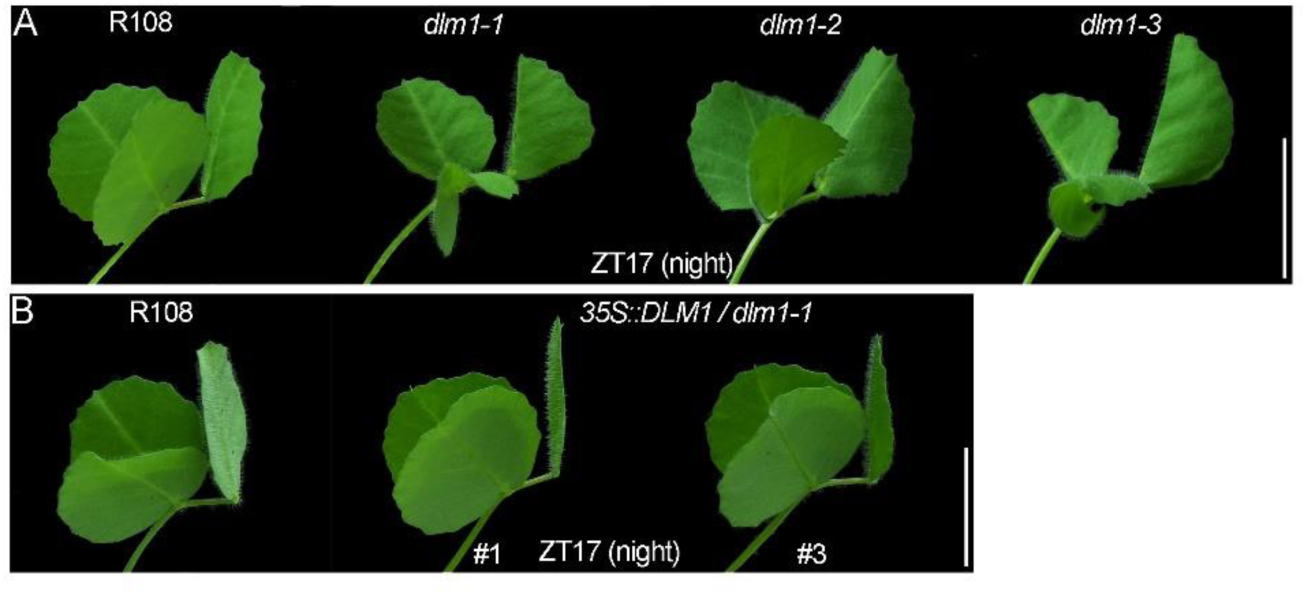
**Phenotype of *dlm1* alleles and genetic complementation at ZT17.** (A) Leaf movement phenotype of R108, *dlm1-1* (NF7737), *dlm1-2* (NF1438) and *dlm1-3* (NF14116) at ZT17 (ZT: zeitgeber) (left to right). Scale bars: 1 cm. (B) The two rescued mutant plants *35S::DLM1*#1 and *35S::DLM1*#3 are shown at ZT17. Scale bars: 1 cm.

**Fig. S6.**
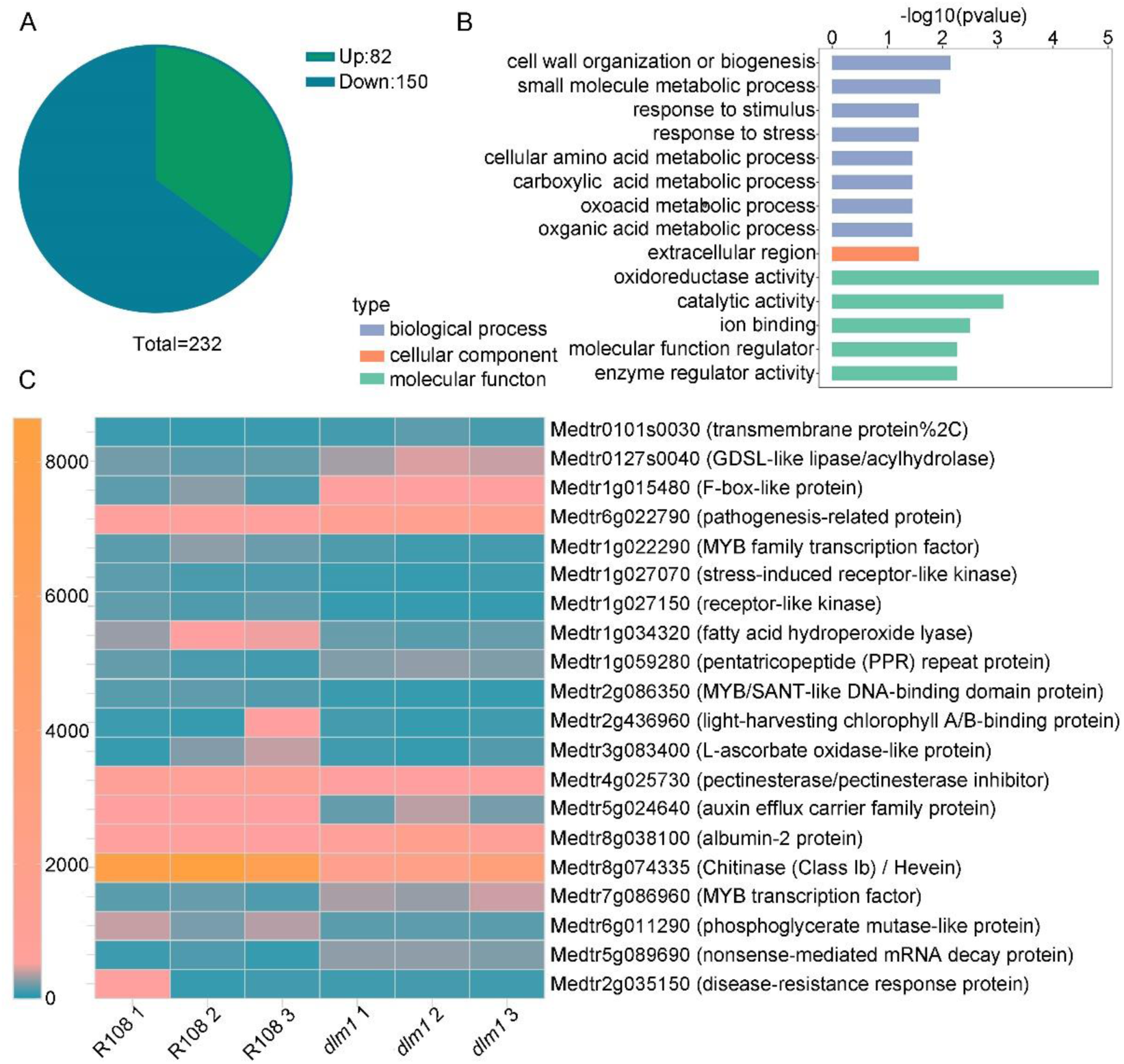
**Transcriptome analysis of pulvinus.** (A) Pie chart analysis of differentially expressed genes. Green represents upregulated genes, blue represents downregulated genes. (B) Gene Ontology (GO) analysis of the DEGs (differentially expressed genes). (C) Heat map showing the representative DEGs with significantly differently expression level.

**Table S1.** *Tnt1* exon insertion sites statistics of *dlm1-1* BC1F2 mutants population by genome resequencing.

| Chromo | Start | End | Locus | Prediction | Annotation |
| --- | --- | --- | --- | --- | --- |
| scf002 | 7581426 | 7581431 | Medtr4g116540 | Heter, Exon | White-brown-complex ABC transporter family protein |
| scf002 | 9400131 | 9400136 | Medtr4g112970 | Heter, Exon | Acyl-ACP-thioesterase B |
| scf003 | 294504 | 294509 | Medtr8g446800 | Heter, Exon | Hypothetical protein |
| scf006 | 4638947 | 4638952 | Medtr6g018770 | Heter, Exon | DUF4378 domain protein |
| scf007 | 2498315 | 2498320 | Medtr1g019960 | Heter, Exon | Alpha/beta-hydrolase family-like protein |
| scf008 | 1210839 | 1210844 | Medtr7g091090 | Heter, Exon | NB-ARC domain disease resistance protein |
| scf008 | 7211644 | 7211649 | Medtr7g103180 | Heter, Exon | Wall-associated receptor kinase-like protein |
| scf009 | 869980 | 869985 | Medtr3g116920 | Heter, Exon | DUF241 domain protein |
| scf010 | 1201595 | 1201600 | Medtr4g057470 | Heter, Exon | DUF1635 family protein |
| scf016 | 10042096 | 10042101 | Medtr6g087000 | Home, Exon | Cyclic nucleotide-binding domain protein |
| scf017 | 3292918 | 3292923 | Medtr2g090580 | Heter, Exon | Early nodulin-like protein |
| scf017 | 6361723 | 6361728 | Medtr2g098180 | Heter, Exon | AP2-like ethylene-responsive transcription factor |
| scf018 | 7764923 | 7764928 | Medtr4g025730 | Heter, Exon | Pectinesterase/pectinesterase inhibitor |
| scf019 | 5802525 | 5802530 | Medtr2g032870 | Heter, Exon | Hypothetical protein |
| scf019 | 6609426 | 6609431 | Medtr2g030460 | Heter, Exon | Stress up-regulated Nod 19 protein |
| scf020 | 1614089 | 1614094 | Medtr2g070410 | Heter, Exon | Caffeoyl-CoA O-methyltransferase |
| scf023 | 1456266 | 1456271 | Medtr3g014530 | Heter, Exon | CC-NBS-LRR resistance protein, putative |
| scf023 | 4904325 | 4904330 | Medtr3g005740 | Heter, Exon | Transmembrane amino acid transporter family protein |
| scf006 | 13169754 | 13169759 | Medtr6g048090 | Homo, 5'UTR | Malectin/receptor-like kinase family protein |

**Table S2.**
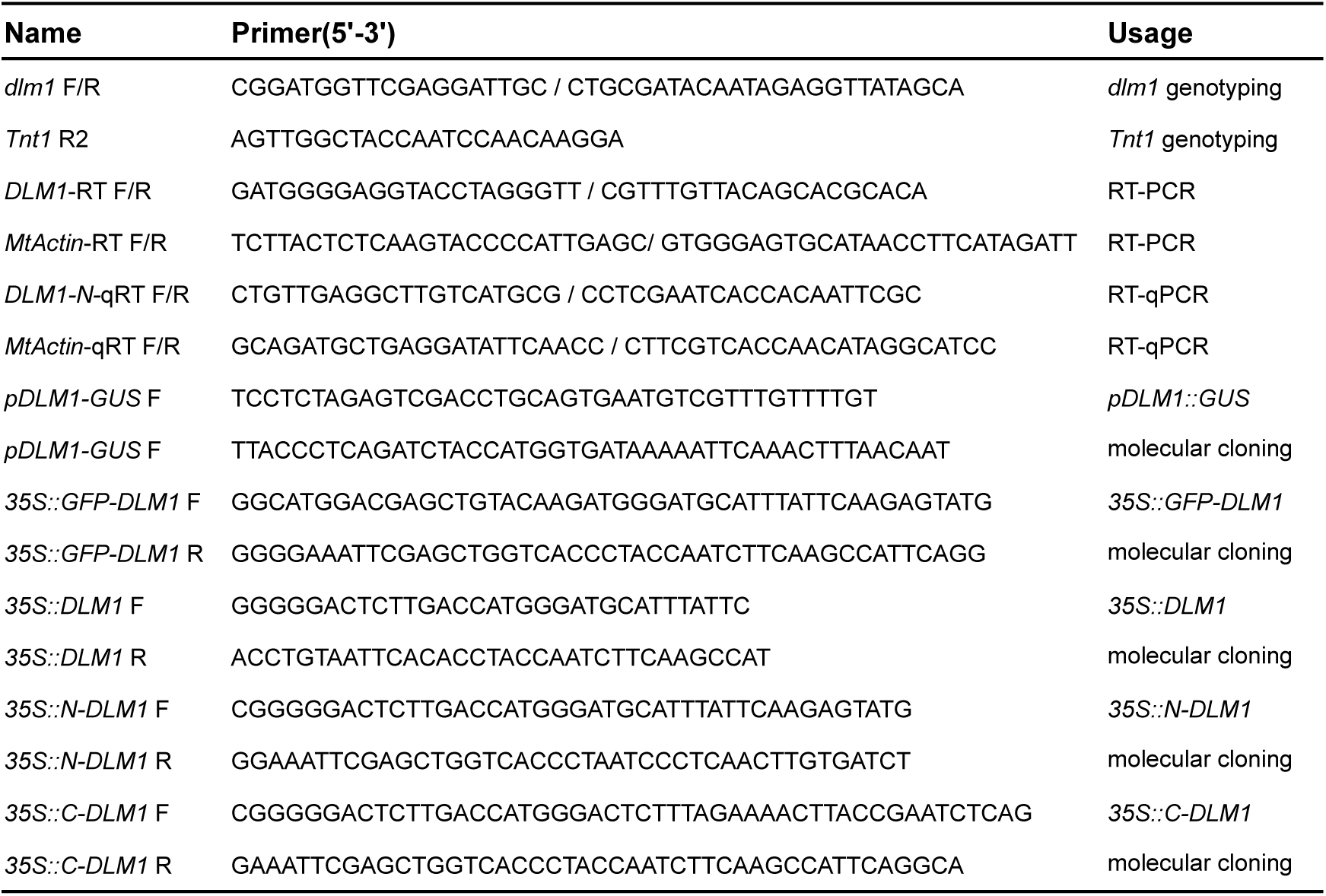
List of primer sequences.

## References

Atamian, H.S., Creux, N.M., Brown, E.A., Garner, A.G., Blackman, B.K., and Harmer, S.L. (2016). Circadian regulation of sunflower heliotropism, floral orientation, and pollinator visits. Science 353, 587–590.

Bai, Q., Yang, W., Qin, G., Zhao, B., He, L., Zhang, X., Zhao, W., Zhou, D., Liu, Y., Liu, Y., et al. (2022). Multidimensional gene regulatory landscape of motor organ pulvinus in the model legume *Medicago truncatula*. International Journal of Molecular Sciences 23(*8*): 4439.

Bhaskara, G.B., Wen, T.-N., Nguyen, T.T., and Verslues, P.E. (2017). Protein Phosphatase 2Cs and *Microtubule-Associated Stress Protein 1* Control Microtubule Stability, Plant Growth, and Drought Response. The Plant Cell 29, 169–191.

Bhaskara, G.B., Wong, M.M., and Verslues, P.E. (2019). The flip side of phospho-signalling: Regulation of protein dephosphorylation and the protein phosphatase 2Cs. Plant Cell Environment 42, 2913–2930.

Chen, J., Moreau, C., Liu, Y., Kawaguchi, M., Hofer, J., Ellis, N., and Chen, R. (2012). Conserved genetic determinant of motor organ identity in *Medicago truncatula* and related legumes. PNAS 109, 11723–11728.

Chen, J., Yu, R., Li, N., Deng, Z., Zhang, X., Zhao, Y., Qu, C., Yuan, Y., Pan, Z., Zhou, Y., et al. (2023). Amyloplast sedimentation repolarizes LAZYs to achieve gravity sensing in plants. Cell 186, 1–15.

Cote, G.G. (1995). Signal Transduction in Leaf Movement. Plant physiol 109, 729–734.

Darwin, C. (1880). The Power of Movement in Plants (London, UK: J. Murray.).

Ding, Y., Lv, J., Shi, Y., Gao, J., Hua, J., Song, C., Gong, Z., and Yang, S. (2018). EGR2 phosphatase regulates OST1 kinase activity and freezing tolerance in *Arabidopsis*. The EMBO Journal 38.

Feng, J., Meyer, C.A., Wang, Q., Liu, J.S., Shirley Liu, X., and Zhang, Y. (2012). GFOLD: a generalized fold change for ranking differentially expressed genes from RNA-seq data. Bioinformatics 28, 2782–2788.

Gao, J., Yang, S., Cheng, W., Fu, Y., Leng, J., Yuan, X., Jiang, N., Ma, J., and Feng, X. (2017). GmILPA1, Encoding an APC8-like Protein, Controls Leaf Petiole Angle in Soybean. Plant Physiol 174, 1167–1176.

Guo, Y., Shi, Y., Wang, Y., Liu, F., Li, Z., Qi, J., Wang, Y., Zhang, J., Yang, S., Wang, Y., et al. (2022). The clade F PP2C phosphatase ZmPP84 negatively regulates drought tolerance by repressing stomatal closure in maize. New Phytologist 237, 1728–1744.

Hagihara, T., Mano, H., Miura, T., Hasebe, M., and Toyota, M. (2022). Calcium-mediated rapid movements defend against herbivorous insects in *Mimosa pudica*. Nature Communications 13: 6412.

Kerk, D., Templeton, G., and Moorhead, G.B.G. (2008). Evolutionary Radiation Pattern of Novel Protein Phosphatases Revealed by Analysis of Protein Data from the Completely Sequenced Genomes of Humans, Green Algae, and Higher Plants. Plant Physiology 146, 323–324.

Kim, H.Y., Cote, G.G., and Crain, R.C. (1993). Potassium Channels in *Samanea saman* Protoplasts Controlled by Phytochrome and the Biological Clock. Science 260, 960–962.

Komatsu, K., Suzuki, N., Kuwamura, M., Nishikawa, Y., Nakatani, M., Ohtawa, H., Takezawa, D., Seki, M., Tanaka, M., Taji, T., et al. (2013). Group A PP2Cs evolved in land plants as key regulators of intrinsic desiccation tolerance. Nature Communications 4.

Komatsu, K., Takezawa, D., and Sakata, Y. (2020). Decoding ABA and osmostress signalling in plants from an evolutionary point of view. Plant Cell Environment 43, 2894–2911.

Kong, Y., Han, L., Liu, X., Wang, H., Wen, L., Yu, X., Xu, X., Kong, F., Fu, C., Mysore, K.S., et al. (2020). The nodulation and nyctinastic leaf movement is orchestrated by clock gene LHY in *Medicago truncatula*. Journal of Integrative Plant Biology 62, 1880–1895.

Kong, Y., Meng, Z., Wang, H., Wang, Y., Zhang, Y., Hong, L., Liu, R., Wang, M., Zhang, J., Han, L., et al. (2021). Brassinosteroid homeostasis is critical for the functionality of the *Medicago truncatula* pulvinus. Plant Physiol 185, 1745–1763.

Liu, X., Zhu, Y., Zhai, H., Cai, H., Ji, W., Luo, X., Li, J., and Bai, X. (2012). AtPP2CG1, a protein phosphatase 2C, positively regulates salt tolerance of Arabidopsis in abscisic acid-dependent manner. Biochemical and Biophysical Research Communications 422, 710–715.

Ma, Y., Szostkiewicz, I., Korte, A., Moes, D., Yang, Y., Christmann, A., and Grill, E. (2009a). Regulators of PP2C phosphatase activity function as abscisic acid sensors. Science 324, 1064–1068.

Ma, Y., Szostkiewicz, I., Korte, A., Moes, D., Yang, Y., Christmann, A., and Grill, E. (2009b). Regulators of PP2C phosphatase activity function as abscisic acid sensors. Science 324, 1064–1068.

Miao, J., Li, X., Li, X., Tan, W., You, A., Wu, S., Tao, Y., Chen, C., Wang, J., Zhang, D., et al. (2020). OsPP2C09, a negative regulatory factor in abscisic acid signalling, plays an essential role in balancing plant growth and drought tolerance in rice. New Phytologist 227, 1417–1433.

Moran, N. (2007). Osmoregulation of leaf motor cells. FEBS Letters 581, 2337–2347.

Moshelion, M., Becker, D., Biela, A., Uehlein, N., Hedrich, R., Otto, B., Levi, H., Moran, N., and Kaldenhoff, R. (2002). Plasma membrane aquaporins in the motor cells of *Samanea saman*: diurnal and circadian regulation. The Plant Cell 14, 727–739.

Nishimura, T., Mori, S., Shikata, H., Nakamura, M., Hashiguchi, Y., Abe, Y., Hagihara, T., Yoshikawa, H.Y., Toyota, M., Higaki, T., et al. (2023). Cell polarity linked to gravity sensing is generated by LZY translocation from statoliths to the plasma membrane. Science 381, 1006–1010

Novak, B., Kapuy, O., Domingo-Sananes, M.R., and Tyson, J.J. (2010). Regulated protein kinases and phosphatases in cell cycle decisions. Current Opinion in Cell Biology 22, 801–808.

Oikawa, T., Ishimaru, Y., Munemasa, S., Takeuchi, Y., Washiyama, K., Hamamoto, S., Yoshikawa, N., Mutara, Y., Uozumi, N., and Ueda, M. (2018). Ion Channels Regulate Nyctinastic Leaf Opening in *Samanea saman*. Current Biology 28, 2230–2238.

Pertea, M., Kim, D., Pertea, G.M., Leek, J.T., and Salzberg, S.L. (2016). Transcript-level expression analysis of RNA-seq experiments with HISAT, StringTie and Ballgown. Nature Protocols 11, 1650–1667.

Procko, C., Radin, I., Hou, C., Richardson, R.A., Haswell, E.S., and Chory, J. (2022). Dynamic calcium signals mediate the feeding response of the carnivorous sundew plant. PNAS 119(30): e2206433119.

Satter, R.L. (1975). Rhythmic and phytochrome-regulated changes in transmembrane potential in *Samanea* pulvini. Nature 255, 408–410.

Satter, R.L., and Galston, A.W. (1971). Potassium Flux a Common Feature of *Albizzia* Leaflet Movement Controlled by Phytochrome or Endogenous Rhythm. Science 174, 518–520.

Scherzer, S., Federle, W., Al-Rasheid, K.A.S., and Hedrich, R. (2019). Venus flytrap trigger hairs are micronewton mechano-sensors that can detect small insect prey. Nature Plants 5, 670–675.

Schweighofer, A., Kazanaviciute, V., Scheikl, E., Teige, M., Doczi, R., Hirt, H., Schwanninger, M., Kant, M., Schuurink, R., Mauch, F., et al. (2007). The PP2C-Type Phosphatase AP2C1, Which Negatively Regulates MPK4 and MPK6, Modulates Innate Immunity, Jasmonic Acid, and Ethylene Levels inArabidopsis. The Plant Cell 19, 2213–2224.

Shi, Y. (2009). Serine/Threonine Phosphatases: Mechanism through Structure. Cell 139, 468–484.

Smékalová, V., Doskočilová, A., Komis, G., and Šamaj, J. (2014). Crosstalk between secondary messengers, hormones and MAPK modules during abiotic stress signalling in plants. Biotechnology Advances 32, 2–11.

Sojka, J., Šamajová, O., and Šamaj, J. (2024). Gene-edited protein kinases and phosphatases in molecular plant breeding. Trends in Plant Science 29, 694–710.

Song, S.-K., and Clark, S.E. (2005). POL and related phosphatases are dosage-sensitive regulators of meristem and organ development in Arabidopsis. Developmental Biology 285, 272–284.

Spartz, A.K., Ren, H., Park, M.Y., Grandt, K.N., Lee, S.H., Murphy, A.S., Sussman, M.R., Overvoorde, P.J., and Gray, W.M. (2014). SAUR Inhibition of PP2C-D Phosphatases Activates Plasma Membrane H+-ATPases to Promote Cell Expansion in Arabidopsis The Plant Cell 26, 2129–2142.

Tadege, M., Wen, J., He, J., Tu, H., Kwak, Y., Eschstruth, A., Cayrel, A., Endre, G., Zhao, P.X., Chabaud, M., et al. (2008). Large-scale insertional mutagenesis using the *Tnt1* retrotransposon in the model legume *Medicago truncatula*. The Plant Journal 54, 335–347.

Yin, P., Ma, Q., Wang, H., Feng, D., Wang, X., Pei, Y., Wen, J., Tadege, M., Niu, L., and Lin, H. (2020). SMALL LEAF AND BUSHY1 controls organ size and lateral branching by modulating the stability of BIG SEEDS1 in Medicago truncatula. New Phytologist 226, 1399–1412.

Yu, L.P., Miller, A.K., and Clark, S.E. (2003). *POLTERGEIST* encodes a protein phosphatase 2C that regulates CLAVATA pathways controlling stem cell identity at *Arabidopsis* shoot and flower meristems. Current Biology 13, 179–188.

Zhao, B., He, L., Jiang, C., Liu, Y., He, H., Bai, Q., Zhou, S., Zheng, X., Wen, J., Mysore, K.S., et al. (2020). *Lateral Leaflet Suppression 1* (*LLS1*), encoding the MtYUCCA1 protein, regulates lateral leaflet development in *Medicago truncatula*. New Phytologist 227, 613–628.

Zhao, W., Bai, Q., Zhao, B., Wu, Q., Wang, C., Liu, Y., Yang, T., Liu, Y., He, H., Du, S., et al. (2021). The geometry of the compound leaf plays a significant role in the leaf movement of *Medicago truncatula* modulated by *mtdwarf4a*. New Phytologist 230, 475–484.

Zhou, C., Han, L., Fu, C., Chai, M., Zhang, W., Li, G., Tang, Y., and Wang, Z.Y. (2012). Identification and characterization of *petiolule-like pulvinus* mutants with abolished nyctinastic leaf movement in the model legume *Medicago truncatula*. New Phytologist 196, 92–100.

Zhou, S., Yang, T., Mao, Y., Liu, Y., Guo, S., Wang, R., Fangyue, G., He, L., Zhao, B., Bai, Q., et al. (2021). The F-box protein MIO1/SLB1 regulates organ size and leaf movement in *Medicago truncatula*. Journal of Experimental Botany 72, 2995–3011.

